# Is the whole the sum of its parts? Neural computation of consumer bundle valuation in humans

**DOI:** 10.1101/2025.04.28.650827

**Authors:** Logan Cross, Ryan Webb, John P. O’Doherty

## Abstract

Humans are often tasked with making decisions about bundles of multiple items and very little is known about how the human brain aggregates, computes and represents value in such cases. We investigated how the human brain evaluates consumer items, both individually and in bundles, and how this activity relates to choice behavior. Participants underwent a deep-fMRI scanning protocol while we elicited behavioral valuations for single and bundled items. Behaviorally, we find that bundle values are sub-additively discounted compared to the sum of individual item values. Neurally, we find that the same distributed network in pre-frontal cortex computes the value of a bundle and its constituent individual items, but the value representation undergoes a normalization that actively re-scales across bundle and single item contexts. These findings suggest that generalized value regions contextually adapt within a valuation hierarchy, as opposed to utilizing an absolute value code.

## Introduction

In daily life, humans must evaluate options that contain multiple components, such as a meal, a bundle of TV channels, an investment fund, or a vacation package. Because decisions over bundles necessitate trade-offs between their distinct components, bundles are the fundamental primitive of economic analysis of consumer choice (Mas-Colell et al., 1995). From a normative standpoint, bundle evaluation can be accomplished via a hierarchical process of valuing the bundle’s constituent items and then aggregating them to value the bundle: the whole is the sum of its parts. This process is formalized in attribute integration theories of value, which propose that the value of a stimulus is constructed by assigning values to each attribute and then aggregating them (Bettman et al., 1998). In the human brain, there is accumulating evidence for partially distinct neural representations of individual features of a stimulus, which are proposed to be integrated to compute an overall stimulus value (Kahnt et al., 2011; Lim et al., 2013; Suzuki et al., 2017; O’Doherty et al., 2021). This evidence is consistent with a large neuroeconomic literature suggesting the existence of a “common currency” value representation in regions of frontal cortex for the objects of choice, including objects with multiple distinct attributes (Levy and Glimcher, 2012b; Bartra et al., 2013; Clithero and Rangel, 2013). However previous research has been inconclusive about the presence of such a unified value signal in bundling (Zyuzin et al., 2022), perhaps due the complexity of valuing and integrating distinct items. In the present study, we examine whether the attribute integration theory of value generalizes to valuing bundles, for which the components that need to be integrated simply represent a different level of abstraction: they are also choice objects. This raises fundamental questions regarding how the value of bundles of items are represented in the brain, and how these representations influence human choice behavior.

Neuroimaging studies have identified specialized cortical areas in lateral orbitofrontal cortex (lOFC) that encode stimulus attributes and exhibit functional connectivity to medial orbitofrontal cortex (mOFC) and ventromedial prefrontal cortex (vmPFC) for value integration (Suzuki et al., 2017; Lim et al., 2013). Furthermore, distinct value representations for categories of goods, such as food and noncomestible consumer items, are separately encoded in posterior regions of the medial orbitofrontal cortex and more anteriorly in the vmPFC, respectively, along with an integrated value signal more dorsally within the vmPFC consistent with a form of “common currency” across categories (McNamee et al., 2013; Clithero and Rangel, 2014). If the features of individual items in a bundle are considered constituent attributes that are combined to compute an integrated bundle value, a similar integrative process can potentially be found for bundles. We might therefore find that values of items and bundles exhibit a hierarchical representation over cortical areas, perhaps continuing the previously-noted lateral-to-medial or ventral-dorsal dissociations. Alternatively, the same region may code for the value of items and bundles, suggesting a more generalized valuation process across different decision contexts. We seek to answer whether there are non-overlapping value representations for bundles that are distinct from the representations of individual items, or whether there is a shared representation used in both contexts. We test these hypotheses by establishing whether spatially distinct brain regions separately correlate with the values of individual items and bundles.

A related question concerns how single-item and bundle value codes are scaled, particularly if items and bundles are evaluated in the same region. Recent behavioral and neuro-physiological evidence suggests that valuation regions in the primate brain implement a relative value code that adapts to the temporal and spatial context of the choice set (Padoa-Schioppa, 2009; Louie et al., 2015a; Rustichini et al., 2017; Khaw et al., 2017; Zimmermann et al., 2018). This compression of the neural code is necessary because neural activity is bounded above and below by physiological limits, and must therefore be scaled to fall within these limits in a manner that maintains sufficient resolution for accurate discrimination. For instance, Cox and Kable (2014) find that the BOLD signal in the brain’s valuation network adapts to the distribution of item stimuli: when the range of items is wider, the responsiveness of the BOLD signal to value attenuates to remain within similar bounds. Behavioral evidence also suggests that a similar re-scaling takes place for attributes (relative to other alternatives) before they are aggregated in the valuation of an item (Dumbalska et al., 2020). However, all of these studies of adaptation have examined this code in the context of valuing single items. When items are combined together, does the neural code simply scale linearly to accommodate the expanded range of possible valuations from the linear combination of items? Or does the neural code adapt across bundle and single item contexts, perhaps by re-scaling each separately?

Given the prior evidence for context-dependent value scaling, we hypothesize that value regions switch between single item and bundle contexts by re-scaling the value code according to the distribution of values in the current context, rather than encoding the absolute value of an option using the same scale across single item and bundle trials. To test this, our experiment has the feature that item and bundle trials are randomly interleaved, so that subjects cannot slowly adapt to one of these contexts. This context-specific normalization would allow for optimal encoding and discrimination of values within each context, aligning with the brain’s tendency to optimize its resources. Context-dependent normalization would also support the remarkable behavioral flexibility observed in human decision-making, allowing individuals to tailor their value representations to the specific demands of the current decision problem, even when choices from multiple contexts are presented in rapid succession. Normalization models including divisive normalization, a canonical cortical computation for encoding relative information (Carandini and Heeger, 2012; Louie et al., 2015b), and range normalization (Rustichini et al., 2017), offer biologically plausible mechanisms for value re-scaling across contexts. Both of these models re-scale valuations and predict that the responsiveness of neural activity will attenuate for bundles relative to single items. For divisive normalization, the responsiveness will attenuate because an item is normalized relative to an additional item (see Methods). For range normalization, it is because the range of valuations increases (Cox and Kable, 2014). We therefore compare the prediction of these normalization codes to alternative relative codes (including a subtractive code and a z-scored code) to shed light on the computational mechanisms underlying the contextual adaptation of the value signal when items are combined together into a bundle.

In our study, we implement an experimental paradigm that is directly comparable to previous research on single item choices (Chib et al., 2009a; Lim et al., 2013; Suzuki et al., 2017). Participants valued single food items, non-comestible consumer goods, and bundles of these same items using an established incentive compatible mechanism (Figure 1). Participants then made choices between these items or bundles and a reference monetary amount in randomly interleaved trials while being scanned with functional MRI (fMRI). To cater for possible variation across participants in the nature of the value codes, and maximize our capacity to probe neural representations of single and bundle values, we adopted an experimental design that was optimized for multivariate representational analyses at the level of individual subjects. We therefore aimed to maximize the number of trials of each type we could obtain per participant by implementing an elongated fMRI experimental design in which each participant performed the task on three separate days of scanning, resulting in over 900 separate trials from each participant. This is many more trials than would typically be found per participant in a conventional fMRI study design. We collected this extensive data in a total of 14 participants, which was also a sufficient sample size to enable group-level inference with a large number of trials.

**Figure 1:**
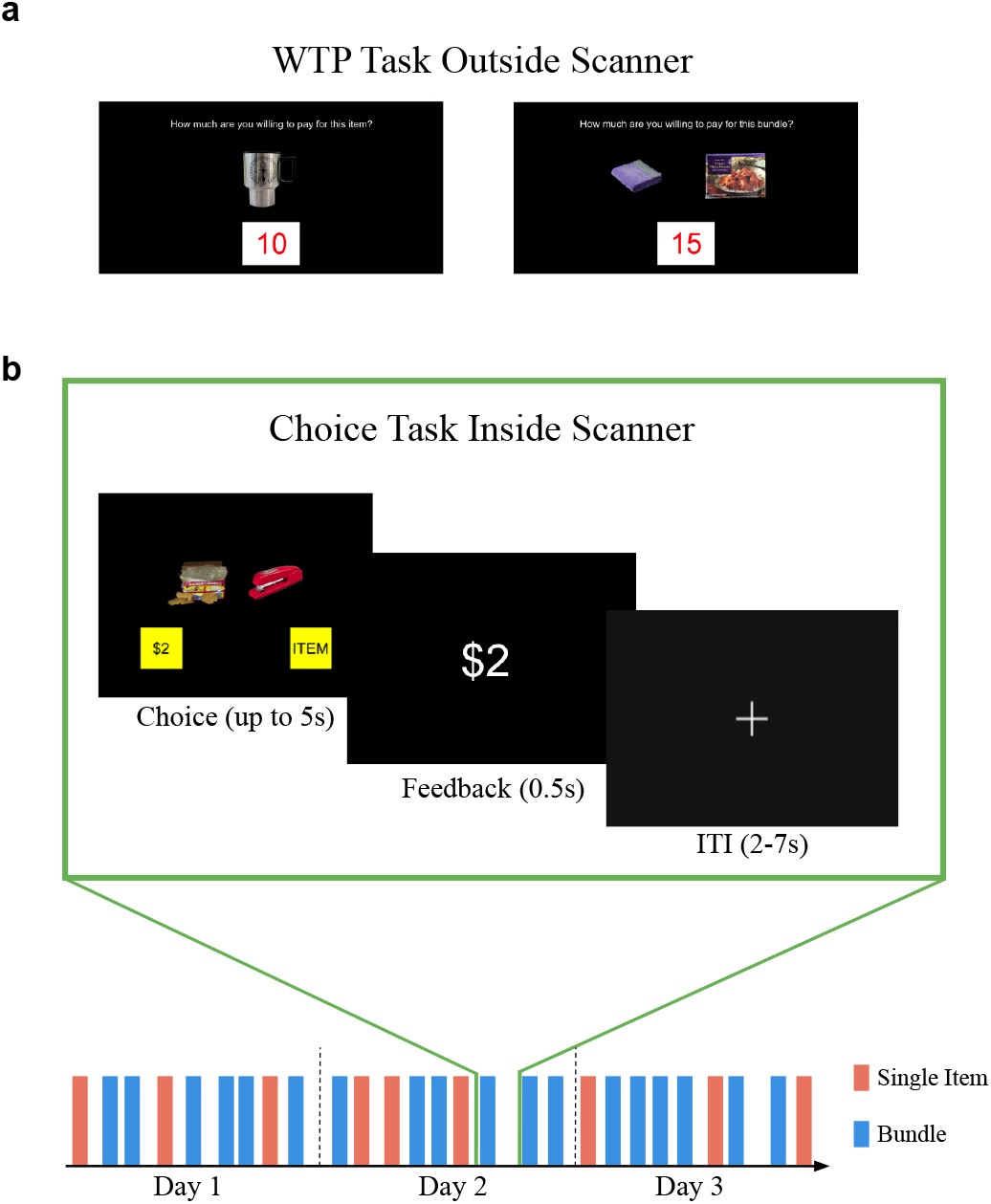
Experimental Design. **a**. WTP task (outside scanner). Participants reported their willingness-to-pay for either items or bundles of two items in a BDM auction. **b**. Choice task (inside the scanner, across three days). During single item trials, a choice was made between an item and a reference monetary amount equal to the median bid of single item trials in the WTP task. Similarly during bundle trials, a choice was made between a bundle and a reference monetary amount equal to the median bid of WTP bundle trials. Single item and bundles trials were randomly interleaved within a run on each day. Participants had up to 5s to make a choice indicated by a right or left button press. The experiment involved 3 days of scanning 5 runs of the task for a total of 15 runs.

## Results

### Bundle value is a sub-additive function of the individual item values

We first examined how the bidding behavior for bundle valuations related to the valuations of the constituent items. Each participant first reported how much they would be willing-to-pay (WTP) for various foods and non-comestible consumer goods (items) and pairs of these items (bundles) (Figure 1). These WTPs are collected using an incentive compatible procedure validated in previous studies, and they therefore provide a measure of an individual’s subjective valuation of each item or bundle (Becker et al., 1964; Chib et al., 2009b; McNamee et al., 2013; Suzuki et al., 2017, see Methods for details). Participants were allocated a budget of $0-$20 and submitted a wide distribution of WTP bids across categories, albeit with most items and bundles valued at small dollar amounts (Figure 2a; mean WTP value for each category: individual item food = $3.34 *±* 3.30 s.d., individual item trinket = $3.30 *±* 3.50 s.d., food bundle = $5.96 *±* 4.44 s.d., trinket bundle = $4.92 *±* 4.81 s.d., mixed bundle $5.394 *±* 4.65 s.d.).

**Figure 2:**
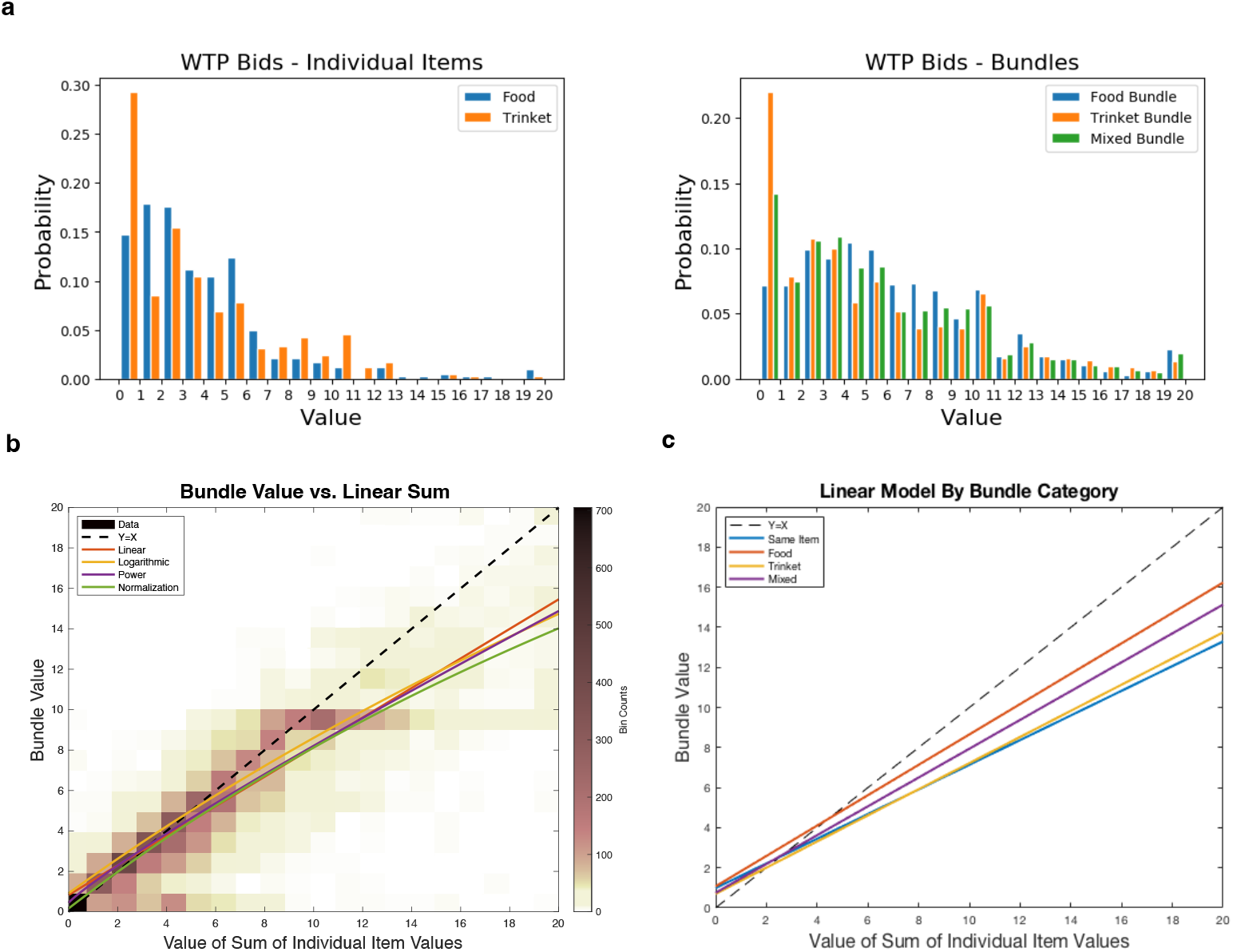
WTP Behavior. **a**. Histograms of WTP bids for individual items and bundles across all subjects. **b**. Density plot of the WTP (value) of a bundle vs. the sum of the values of the constituent items in a bundle (across all subjects). The diagonal represents where bundle values equal the sum of the constituent item values. The model fits from four models are plotted over the density plot. All models conclude that bundle value is a subadditive function of the individual item values, as they extend below the diagonal. **c**. Linear model fits stratified by bundle type.

In Figure 2, we plot the density of bundle valuations relative to the sum of the individual item valuations. While the bundle values tend to increase monotonically with the sum of the individual item values, as might be expected, there is substantial density below the diagonal suggesting that bundle value is a sub-additive function of the individual item values. To test this hypothesis, we modeled the value for a bundle of items, *v*_*i,j*_, as a function of the constituent item values, *v*_*i*_ and *v*_*j*_. The theory of attribute integration specifies the value of a bundle as a linear combination of values of the individual items (*v*_*i,j*_ = *β*_0_ + *β*_1_*v*_*i*_ + *β*_2_*v*_*j*_). Using a linear mixed effects regression, we found that both parameter estimates for the individual items were significantly less than one 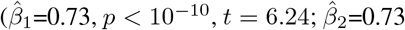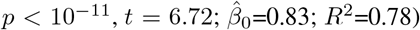, consistent with the hypothesis that bundle value is a sub-additive function of the individual item values (Figure 2b). We additionally tested three non-linear mixed-effects models which allow for concave functions of the combined items (a power model, a logarithmic model, and a divisive normalization model, see Methods). The best-performing model was a divisive normalization model which normalizes the bundle components relative to each other (Δ*BIC* = 182). The power model and logarithmic model, which allow for parametric non-linear functions on the valuations, also fit the data better than the linear model (Δ*BIC* = 129, Δ*BIC* = 65). The parameters of all of these non-linear models are estimated to yield concave value representations (Figure 2b), thus participants discount their subjective value of a bundle more as the individual item values increase.

We also analyzed how the relation between bundle and item value varies depending on the category of the bundle. The linear model fit for each type of bundle is shown in Figure 2c. The slope is smallest for bundles that are duplicates of the same item (*β*_1_ = 0.61*±*0.01, *β*_2_ = 0.61*±*0.01, *β*_0_ = 0.98*±*0.13), which is consistent with the economic principle that the utility of an item decreases with each additional unit (diminishing marginal utility). Moreover, food and mixed bundles have larger slopes than same-item bundles and trinket bundles (Food: *β*_1_ = 0.74*±*0.04, *β*_2_ = 0.77*±*0.04, *β*_0_ = 1.07*±*0.26; Mixed: *β*_*Food*_ = 0.71*±*0.05, *β*_*T rinket*_ = 0.71*±*0.06, *β*_0_ = 0.77*±*0.26; Trinket: *β*_1_ = 0.67*±*0.06, *β*_2_ = 0.63*±*0.06, *β*_0_ = 0.70*±*0.19).

### Neural representation of subjective value and choice

We next investigated how subjective value and choice are encoded in the brain during the choice task. After the WTP task, participants were scanned using functional MRI (fMRI) while they performed a choice task with the same items and bundles. Three participants were scanned with a high-resolution fMRI protocol (voxel size = 1.5mm isotropic) designed to record from medial prefrontal cortex (mPFC) regions with high fidelity (Figure S1). The remaining eleven participants were scanned with a standard wholebrain protocol (2.5 mm isotropic). On each trial in the choice task, a participant made a choice between an item (or a bundle) vs a reference monetary amount equal to their median WTP bid on that category (Figure 1b). This ensured that participants would value the item/bundle in isolation and choose the item/bundle about half the time. We observed a slight bias towards choosing the money on individual item trials (Figure S3).

#### Univariate Analysis

To examine whole-brain correlates of bundle and single item valuation, we first constructed a univariate general linear model (GLM) that included regressors for a stimulus WTP rating time-locked to the onset of the trial, and a regressor for the choice made on that trial time-locked to the choice (in addition to a regressor for trial type and other covariates of no interest; see Methods). Several regions were more active when participants chose an item or bundle vs. when they chose the reference monetary amount, including clusters in dmPFC areas such as superior frontal gyrus (SFG), anterior cingulate gyrus (ACC), vmPFC, and the angular gyrus (*p <* 0.001 FDR corrected cluster-level, Figure 3a). The three participants that underwent the high-resolution scanning protocol are absent from this specific whole-brain analysis because that protocol was focused on ventral and frontal areas (with a limited field of view) with no coverage in dorsal frontal areas. By including the regressor for the WTP of the item/bundle shown in a trial in the same GLM, we were also able to isolate subjective value signals independent of choice. After cluster-level false discovery rate correction (*p <* 0.001, FDR), one large cluster in the anterior portion of vmPFC and frontal pole showed a positive correlation to subjective value (Figure 3b). These results are consistent with previous findings of a general value signal (Bartra et al., 2013; Clithero and Rangel, 2014; Levy and Glimcher, 2012a) and set the stage for further examination of how the representations in these regions are modulated over single item and bundle trials.

**Figure 3:**
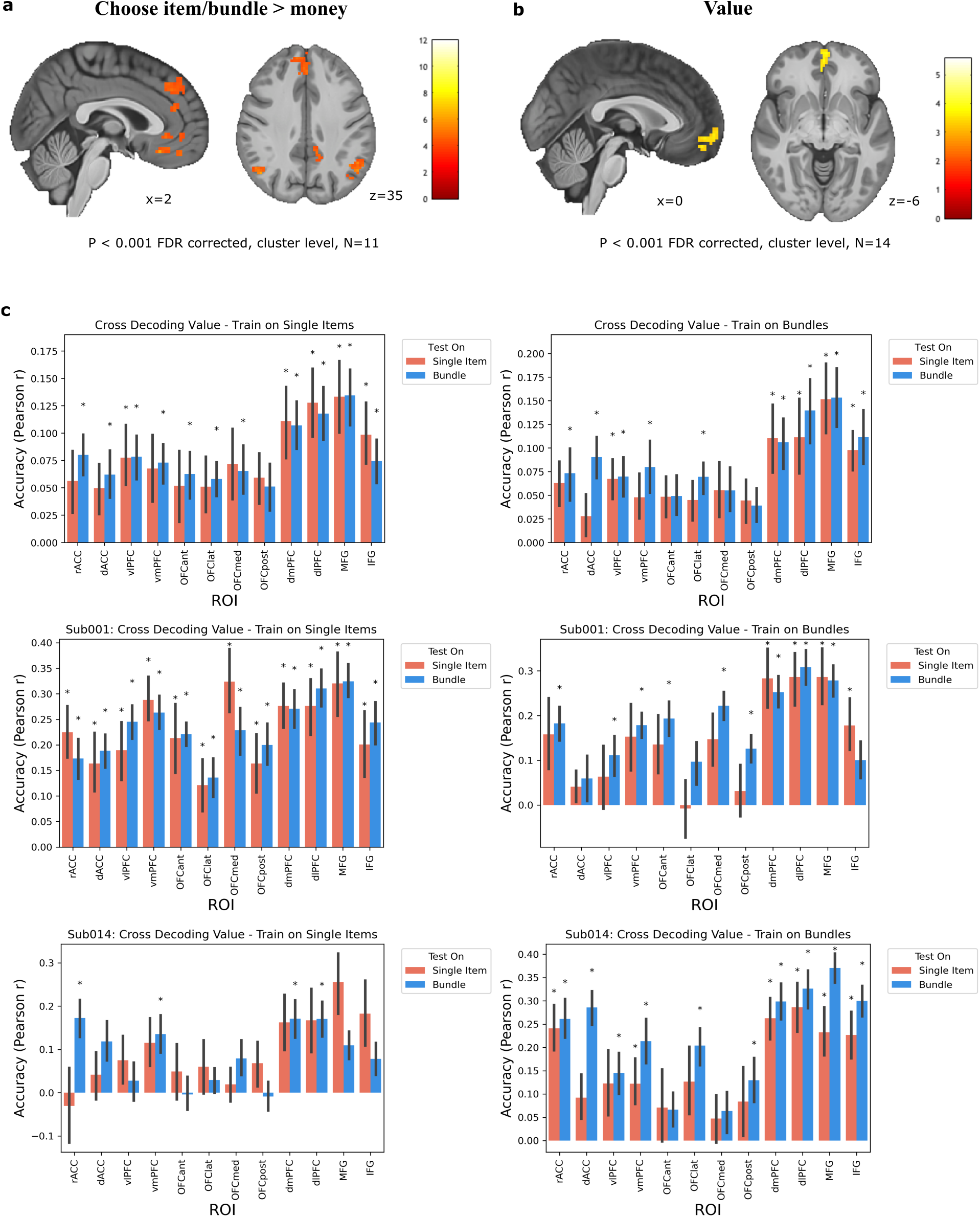
Neural representation of subjective value and choice. **a**. Areas more active when the item or bundle was chosen than when the reference monetary amount was chosen. Significant clusters in dmPFC/SFG, ACC, vmPFC, and angular gyrus (N=11). Clusters are defined by a *p <* 0.001 uncorrected cluster forming threshold followed by *p <* 0.001 FDR cluster-level correction. **b**. Neural correlates of the value of the item/bundle presented. Significant cluster in anterior vmPFC (N=14). Clusters are defined by a *p <* 0.001 uncorrected cluster forming threshold followed by *p <* 0.001 FDR cluster-level correction. **c**. MVPA cross decoding analysis results (N=14). Ridge regression decoders were trained on samples from a condition and tested on held-out samples from either condition (Left^1^: ^0^trained on single items, Right: trained on bundle trials. Top row: Group result. Bottom rows: the first and the last subject). Asterisks * represent significant prediction accuracies on a test partition (two-sided one-sample Wilcoxon signed rank test *p <* 0.05 and FDR-corrected for multiple comparisons *q* = 0.05). At the group level, there were no significant paired differences in prediction accuracies between test conditions for any ROI. Error bars reflect SE across participants.

We next investigated whether anatomically distinct brain regions represent the value of individual items and the value of bundles. One possibility is that additional regions are recruited when computing the value of bundles, since this involves a hierarchical process of valuing the individual items and then integrating them to evaluate the bundle. To examine whether the brain recruits additional circuitry when evaluating a bundle versus valuing a single item, we tested for the interaction of value and trial type in the previously described GLM (contrasts: bundle value *>* single item value and single item value *>* bundle value). At the group level, no clusters survived correction for either comparison. Therefore we do not find evidence for a topography of separable single item value and bundle value codes at the univariate level (Figure S3).

#### Multi-variate Analysis

We next aimed to determine whether the value of bundled items might be represented differently from single items at a more distributed level. We utilized multivariate pattern analysis (MVPA) in order to detect such distributed codes. Using all 14 subjects, we implemented decoding analyses across conditions to test for the existence of distinct bundle value codes, single item value codes, or general value regions. A ridge regression decoder was trained on distributed voxel patterns in several regions of interest (ROIs) in PFC. The decoder was trained on samples of the valuations from one trial type (e.g. single-item trials) and then tested separately on a held out run from either trial type (i.e. a leave-one-run-out cross-validation). Decoder performance was assessed with a Pearson correlation between the predicted values and the true values in the test set. To test the hypothesis of a condition-independent general value region, we analyzed if the decoder could predict the valuations of samples drawn from the other condition. This would identify general value regions that utilize the same distributed code when computing the value of both single items and bundles. To test for condition-dependent value regions, we analyzed if the decoder could only predict samples within the condition it was trained on, while failing to predict samples from the other condition. This would identify distinct single-item value regions or bundle value regions.

We found that value could be predicted at the group level on all four types of train/test splits (‘train and test on single items,’ ‘train and test on bundles,’ ‘train on single items/test on bundles,’ ‘train on bundles/test on single items’) in vlPFC, dmPFC, dlPFC, MFG, and IFG (Figure 3c, two-sided one-sample Wilcoxon signed rank test P *<* 0.05 and FDR-corrected for multiple comparisons *q* = 0.05). Decoding value across both conditions in these regions suggests that they possess distributed general value codes that are independent of condition. Additionally, value was decoded above chance for the ‘train on single items/test on bundles’ partition in rACC, dACC, vmPFC, anterior OFC, lateral OFC, and medial OFC, providing some evidence for generalized value codes in these regions as well (although prediction accuracies were not significant for the reverse partition ‘train on bundles/test on single items’).

In order to depict how value decoding varies across individuals, individual subject results are also shown for two representative subjects in Figure 3c, and the remaining subjects in Supplementary Figure 4. To identify a condition-dependent value region, we assessed if a region’s decoder only had significant prediction accuracy when trained and tested on a single condition (i.e. only significant for ‘train and test on single items’ but not for the other three partitions). No region met this criteria at the group level. Additionally, there were no significant differences in a decoder’s prediction accuracy between conditions for any ROI (two-sided two-sample Wilcoxon signed rank test *p <* 0.05 and FDR-corrected for multiple comparisons *q* = 0.05). For example, although there is a 0.06 difference in pearson correlation between ‘train on bundles/test on single items’ and ‘train and test on bundles’ for dACC, this difference is not significant after correcting for multiple comparisons (*p <* 0.426). Thus, the cross decoding analyses do not yield any evidence for the existence of a distributed item and/or bundle-specific value code in any ROI. Instead, these findings are consistent with a distributed general value code for both bundles and single items.

**Figure 4:**
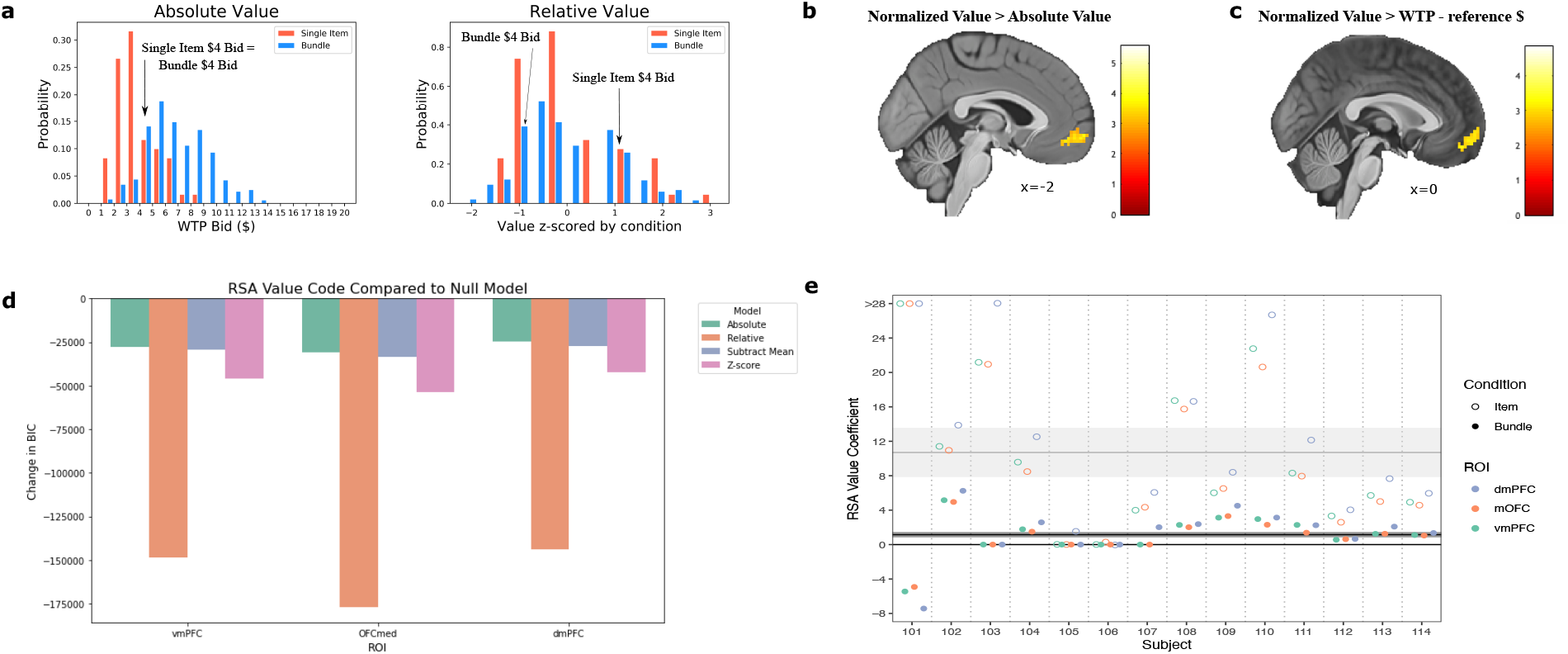
Normalization of the value code. **a**. A depiction of the distributions of value by condition with absolute value and relative value codes. An absolute value code (left) represents value according to the participants’ WTP bids, an incentive compatible measure of the subjective value of an item or bundle. A relative value code normalizes value by condition, plotted here after z-scoring individual item values and bundle values separately for an example subject (right). A relative code adapts to the context of the task and puts values from different distributions on the same scale. **b**. Contrast of normalized value *>* absolute value. A cluster in vmPFC is better fit by the normalized value model, indicating that the value representation in this region is normalized. Clusters are defined by a *p <* 0.001 uncorrected cluster forming threshold followed by *p <* 0.001 FDR cluster-level correction (N=14). **c**. Contrast of normalized value *>* value difference, with a cluster in vmPFC similar to the previous pane. Value difference is the WTP bid $ amount - the reference $ amount. Results are *p <* 0.005 FDR cluster-level corrected (*p <* 0.001 uncorrected cluster forming threshold) (N=14). **d**. Model comparison (change in BIC) of different representations of value compared to the null model (wih no value component but including nuisance controls). Models are fit via representational similarity analysis (RSA). More negative values imply a better model fit — thereby implicating the relative value model as the best account for the neural code in each region (N=14). **e**. RSA Value coefficients from the Relative model, by subject. Value coefficients are attenuated for Bundle trials relative to the Item trials, as predicted by normalization models. Horizontal lines reflect the average coefficient the standard error across all subjects and ROIs.

### Normalization of the value code

If the brain’s value regions represent value in a manner that generalizes across individual items and bundles of those items, a subsequent question emerges: how does the value code adaptively normalize to trial type? The common currency hypothesis suggests that options are encoded with the same value scale so they can be compared (Padoa-Schioppa and Assad, 2008; Chib et al., 2009b; Lin et al., 2012; Levy and Glimcher, 2011), thereby value regions may code for items and bundles on the same range that scales with their WTP ratings. However, given the biological constraints of neurons, in order to evaluate decisions on drastically different value scales with the same valuation system (e.g. decisions about which house to buy or which entree to buy), the brain must also adaptively normalize the value code to the distribution of values available in a decision-making context (Louie et al., 2015a). Since the distribution of bundle values is skewed toward larger values compared to the distribution of single item values (Figure 2a), the neural code may normalize to the context of these different trials. A value region could therefore encode a $4 rated item very differently than a $4 rated bundle if options are appraised based on their utility relative to the other options within a condition (Figure 4a). Previous work has established relative value coding empirically in experiments where a different distribution of values is presented in a block design (Padoa-Schioppa, 2009; Louie et al., 2011; Khaw et al., 2017; Zimmermann et al., 2018), presumably because the value code adapts gradually over time to match the distribution in each block. But crucially in our experiment, bundle trials and single item trials are randomly intermixed within a block. Therefore any adaptation, if present, must be instantaneous rather than slowly learned.

To test whether the neural representation codes for value in absolute or relative terms, we constructed GLMs with different value regressors in all subjects. One GLM included a regressor for value according to the WTP bid in dollars for the item or bundle displayed in that trial, while in a separate GLM value was normalized (with a z-score) by trial type. Value contrasts were then compared at the second level (normalized value *>* absolute value and absolute value *>* normalized value). A significant cluster in vmPFC emerged for the normalized value *>* absolute value comparison (peak voxel: x=-2, y=62, z=-7; t_13_=5.75; *p <* 0.001 uncorrected cluster forming threshold followed by *p <* 0.001 FDR cluster-level correction), suggesting that the neural representation of value in vmPFC is normalized. No significant clusters emerged after correction for the opposite analysis (absolute value *>* normalized value). An alternative explanation for this result is that vmPFC is computing the difference in value between the item/bundle presented in a trial and the reference monetary amount it is choosing against. This subtractive normalization computation would also produce a relative value code, but is a simpler form of adaptation that does not place the single item value and bundle value distributions on the same scale. We therefore constructed another GLM with a modified value regressor equal to the WTP bid minus the reference monetary amount, and the resulting value contrast was tested against the fully normalized value contrast. Again, a significant cluster in vmPFC resulted from the normalized value *>* value difference contrast (peak voxel: x=0, y=52, z=-14; t_13_=4.83; *p <* 0.001 uncorrected cluster forming threshold followed by *p <* 0.005 FDR cluster-level correction), and the opposite contrast yielded no significant clusters. These results demonstrate that at least at the univariate level, vmPFC normalizes the value code by condition.

To examine a relative value representations at the multivariate level, we performed representational similarity analysis (RSA) in all subjects. RSA provides a data-driven method to examine the structure of neural representations by constructing dissimilarity matrices (DSMs) according to how the multidimensional voxel space changes from trial-to-trial or from condition-to-condition (Kriegeskorte et al., 2008). These data-driven DSMs can then be compared to model DSMs that encode how the features of the task, such as value, evolve from trial to trial. We built trial-by-trial DSMs for every ROI and correlated them to model DSMs for four potential value representations: absolute value, subtractive normalization (subtracting the mean value of a stimulus in a category), z-score normalization, and a relative normalization model which allows the value coefficient to differ over trial type (item and bundle). In the latter model, this value coefficient will reflect the responsiveness of the neural code to value. Both divisive normalization and range normalization predict that this responsiveness should attenuate on bundle trials relative to single-item trials. For divisive normalization, the responsiveness will attenuate because an item is normalized relative to an additional item (see Methods). For range normalization, it is because the range of valuations increases. Therefore an attenuation of the value coefficient would be consistent with a relative normalization model, but inconsistent with an absolute value model.

We compared all of the above models to a null model which only contains nuisance regressors. As shown in Figure 4d, the relative normalization model of value provides a much better fit in three value ROIs at the group level, vmPFC, OFCmed, and dmPFC. Additionally, over all subjects, the average value coefficient is significantly lower in bundles than for single items in all three of these areas (*t* = *−*3.359, *p <* 0.001). This suggests that the neural geometry in these value regions is most reflective of a relative value code in which the responsiveness of neural activity is attenuated in bundle trials.

## Discussion

In real-world decision-making, consumers often have to make choices between options that are each made up of multiple goods. However, it is unknown how the human brain constructs the value of bundles of multiple items. To investigate this question, we used a BDM auction procedure to elicit subjective values for food items, non-comestible consumer goods, and bundles of these items. Then participants made choices with these items and bundles during fMRI scanning.

We found that bundle value is computed with a sub-additive function of the individual item values, therefore bundles are systematically discounted in relationship to the sum of the individual item values. Concave non-linear models capture this behavior, and this concavity suggests that participants discount the subjective value of a bundle more as the values of its constituent items increase. Our best-performing behavioral model normalizes the value of the items relative to their sum; however, other nonlinear models also do a comparable job of capturing this sub-additive behavior.

At the neural level, we tested for absolute or relative value signals, across single item and bundle choices, at the univariate and multivariate levels. Previous research has identified a spatial topography in OFC where food value was represented in posterior mOFC and value codes for consumer goods were represented in anterior mOFC (McNamee et al., 2013). In the present study, we investigated whether a spatial topography would also exist in relation to the valuation of single items versus bundled items. One possibility that we tested is that bundle items, being more complex in the sense of being composed of multiple items as features, would be represented in a distinct region of the vmPFC compared to the value of single items. However, we found no evidence for separate single item value regions or bundle value regions, or for a topography of value complexity. An anterior portion of vmPFC showed a correlation to subjective value across both individual item trials and bundle trials. This result is consistent with a large body of literature finding category-independent value signals in vmPFC (Bartra et al., 2013), however evidence for this signal in bundle valuation has been elusive (Zyuzin et al., 2022). Given our finding that the BOLD signal responsiveness to value is attenuated in bundle trials, this may have required higher-powered analyses to observe. In a cross-decoding analysis similar to methods previously used to identify category-dependent and category-independent value codes (McNamee et al., 2013), distributed value codes were revealed throughout PFC, including in dmPFC, vlPFC, and MFG for both bundles and single items, however we found that none of these regions differentially coded for one or the other. This suggests that pre-frontal cortex represents a generalized value signal across a valuation hierarchy of items and bundles.

While we did not find evidence for any spatial topography with regard to the representation of the value of single and bundled items, we did find evidence to suggest that value is encoded in these regions in a relative as opposed to absolute manner. Specifically, we found that a form of normalization of value representation is occurring in these areas, such that the value representation is attenuated on bundle trials relative to single item trials. Using representational similarity analysis, we compared neural activity against different forms of normalized and non-normalized value signals. We find that a scale normalized value code provides the best account of the neural data compared to other forms of non-normalized value signals. This results is consistent with both divisive and range normalization models, which predict an attenuation of the value representation for bundles.

Our findings have implications for the common currency hypothesis of value coding (McNamee et al., 2013; Clithero and Rangel, 2014). Partly consistent with the notion of a common currency, we found evidence that the value regions we identified in our study encode value in a general way across both single-items and bundles, and both within and across categories of goods. While our results show that value is represented by the same distributed voxel pattern, we find that this value code does normalize across contexts. Thus, it is clear that our findings do not support a context-invariant common currency representation. Instead, the brain appears to be utilizing a value representation that is scaled by context (Louie et al., 2011; Padoa-Schioppa and Assad, 2008). While we highlight divisive or range normalization as possible mechanisms for explaining this attenuation in bundle trials, our experimental designs cannot distinguish between them (or exclude other possible normalization mechanisms). Future studies which alter the number and composition of bundles are needed to do so (Daviet and Webb, 2023, e.g.).

Our behavioral results are compatible with the expectation most consumers have that bundles of multiple goods are usually discounted in comparison to the total price of the constituent products purchased separately (Thorne, 2004; Nagle and Holden, 1987). Value meals at fast food restaurants, snack variety packs, vacation packages, and season tickets for sports teams all represent discounted bundles in the real-world marketplace. However, the incentives of buyers and sellers in the marketplace are usually different than in our experimental setup. Sellers typically offer bundles at a discount to strategically ensure that high-willingness to pay customers pay a higher price, as in discounting an item when it is purchased in bulk (i.e. second degree price discrimination). Or alternatively, they increase the probability of a buyer purchasing an additional good they would not have purchased otherwise. For example, a customer at a fast-food restaurant will often have the option to buy a hamburger and fries separately, so a bundle of the two needs to be discounted in order to become an attractive package. In our experiment, each item and bundle is bid on separately and a BDM auction trial is selected at random, so when evaluating a bundle in the experiment, the option to purchase them separately is not available. Therefore, the results of our experiment suggest that bundle discounting may be a more general phenomenon of multi-item valuation and not just the result of equilibrium pricing strategies in a market. This pattern is consistent with the law of diminishing marginal utility, which holds that an additional unit of a good has a lower value. In our experiment, bundles of the same item were discounted slightly more than the other bundles.

Microeconomic theory also accounts for the possibility that bundles of goods are substitutes or complements (Varian, 2014). For example, tea and coffee could be viewed as substitutes because they both offer a caffeine boost. A consumer typically does not buy both at a cafe, so their utility bundled together is likely less than the sum of their independent utilities. Complements are products which are typically bought and used together, such as pasta and pasta sauce. Other examples, like left and right shoes, can exhibit super-additivity, where the utility of one shoe by itself is lower than half of the value of both shoes together. However, in economic theory, these concepts are typically defined in terms of a consumer’s response to price changes, which are not present in our experiment. Moreover, the concepts of diminishing marginal utility and substitution/complementarity are not intrinsically linked theoretically: it is possible to define a utility function over perfectly substitutable goods which also exhibits diminishing, constant, or increasing marginal utility. Though we did not construct our experiment to analyze substitution or complementarity due to price changes, it is possible that the sub-additivity we observe in behavior may arise from our participants evaluating a large majority of the stimuli as substitutes with diminishing marginal utility. Investigating how the brain evaluates and represents the value of bundles containing such items, and how their attributes are traded off against each other, is an interesting avenue that we hope to leverage for future brain research. In particular, a study design which directly manipulates the substitutability of items, rather than just randomizing them into bundles, could examine how these neural representations might differ.

To conclude, in the present study we show that the brain represents the value of bundles in the same brain areas that encode the value of single items. We found no evidence for a topographical arrangement of value representations within the ventromedial prefrontal cortex or elsewhere, when comparing single item and bundled item value signals, suggesting that these value signals are not hierarchically structured at least with respect to neuroanatomical organization. However, the responsiveness of brain activity within those brain regions is attenuated for bundles relative to single items. Our findings thus indicate that value representations for complex combinations of stimuli utilize similar neural representations, but these are scaled differently depending on their context, as opposed to utilizing an absolute value code.

## Acknowledgements

This work was supported by a National Institute of Mental Health Caltech Conte Center grant project on the neurobiology of social decision-making (P50MH094258) to JPOD, a National Institute on Drug Abuse (R01DA040011) grant to JPOD and LC, and a SSHRC Insight Development Grant (430-2019-00246) to RW. We would like to thank Whitney Griggs for assisting with data collection.

## Methods

### Participants

Participants (N=14) were recruited from the general population through the Caltech Brain Research Participant System (7 females, 7 males, 24.9 *±* 3.74 years, mean *±* s.d.). They did not have any food allergies and were not dieting at the time of the experiment. They were given a participation fee of $40 ($20 per hour), in addition to receiving monetary, food, and non-comestible consumer goods as rewards depending on their choices in the experiment. Each subject gave their informed consent, and the study was approved by the Institutional Review Board of the California Institute of Technology.

### Stimuli

Across participants, 70 food items and 40 non-comestible consumer goods were used as stimuli in the experiment. Food items included fruits, snacks, and mains (including microwaveable meals) that are available at local grocery stores. Consumer goods included a diverse array of items under $40 in price, including cell phone chargers, kitchen items, Caltech memorabilia, and books. Many of these items have been used in previous studies (Suzuki et al., 2017).

The full list of items can be found in the Supplementary Table 1.

**Table 1.** Behavioral models of bundle value. R^2^ and BIC scores reflect fit across participants. Each model fits random effects parameters per participant. See Methods for model details.

| Model | R<sup>2</sup> | BIC |
| --- | --- | --- |
| Linear | 0.7771 | 39151 |
| Polynomial | 0.7722 | 39510 |
| Power | 0.7797 | 39049 |
| Logarithmic | 0.7777 | 39097 |

### Experimental tasks

Participants performed the experiment in three sessions on three separate days in order to maximize the amount of fMRI data within-subject. On each day, participants first performed a willingness-to-pay (WTP) task outside the scanner, then performed the choice task in the scanner where they were asked to choose between an item or bundle of two items versus a reference monetary amount. Participants were asked to refrain from eating 4 hours before the experiment in order to ensure that food items were valuable to them. Compliance was confirmed by self-reports, that asked whether a participant complied, in addition to hunger ratings.

Each participant performed the willingness-to-pay and choice tasks on a total of 40 unique items presented throughout the experiment. On a given day, the participant was presented 20 items, composed of 10 food items and 10 non-food items. 10 of these items were presented in all three days, and 10 new items were introduced every day. To construct bundles, every item was paired with each other, including pairs of the same items. Thus, 210 bundles were included each day (20 choose 2 = 190 + 20 pairs of the same item). On the final day of the experiment after the choice task, outside the scanner participants rated how familiar they were with each of the 40 items. For each item, the participant indicated their familiarity with the item on a continuous scale from ‘not at all’ to ‘very much’ by moving a red pointer, with no time constraint as described previously (Suzuki et al., 2017).

### WTP Task (outside the MRI scanner)

Participants completed an un-timed BDM auction task to measure their willingness-to-pay for items and bundles, with a procedure similar to previous studies (Chib et al., 2009b; McNamee et al., 2013; Suzuki et al., 2017). The BDM auction is a reliable incentive-compatible method to elicit subjective values for items (Becker et al., 1964). Participants were endowed with a $20 budget in cash, and instructed that they can use this cash to purchase items from our laboratory store (and keep the money they do not use). In each trial, an item or bundle was shown and the participant was asked to type in how much they would be willing to pay from $0-$20 for that item/bundle (Figure 1). Participants first bid on the individual items (20 each day) and then bid on the bundles (210 each day).

Each trial was to be treated independently, as a random trial from the entire experiment was selected at the end of the experiment. If the selected trial was from the WTP task, the participants’ bid on that trial was then compared against a randomly generated price (uniform probability from $0-$20), and if their bid was greater than or equal to that price, they received the item(s) and paid the corresponding price with their $20 budget. If their bid was less than the price, they did not receive the item(s), and they did not have to pay anything. Participants were explicitly instructed about this auction procedure, and about how the optimal strategy is to bid their true subjective value for a given item/bundle. With a questionnaire, we confirmed that participants understood the mechanism of the auction.

### Choice Task (inside the MRI scanner)

In the scanner, participants made choices involving the items and bundles previously bid on in the WTP task. Each trial involved a binary choice between an item or bundle and a reference monetary amount (Figure 3.1b). The reference monetary amount was equal to the participant’s median WTP bid for that category (individual items were chosen against the median bid on individual items and bundles were chosen against the median bid on bundles). This ensured that the participants would choose the reference monetary amount about half the time and choose the item or bundle half the time. For a trial, the stimulus appeared in the middle of the screen, and the word ‘ITEM’ appeared on the bottom left or right with equal probability while the reference monetary amount in ‘$X’ appeared on the other side. Participants selected their choices with a button box, with the leftmost button indicating choosing the left option and vice versa for the rightmost button. Participants had 5 seconds to make a decision, after which their choice was presented on the screen for 0.5 seconds and followed by a jittered intertrial interval (ITI phase, 2–7s).

On each of the three days, participants were scanned for five runs. Each run included 62 trials, where each of the 20 individual items was presented once per run, and each of the 210 bundles in a day was presented once on that day. On day 1, anatomical scans were collected after the choice task.

### fMRI Data Acquisition

The fMRI data was acquired on a Siemens Prisma 3T scanner at the Caltech Brain Imaging Center (Pasadena, CA) with a 32-channel radio frequency coil. At the end of the first day of scanning, T1 and T2 weighted anatomical high-resolution scans were collected with 0.9mm isotropic resolution.

#### High resolution data

A high-resolution partial volume slab was collected in three participants with a 1.5mm isotropic voxel size (Figure S1) and the following parameters: multiband acceleration = 4, 64 slices, TR = 1100ms, TE = 26ms, flip angle = 63°, FOV= 192mm x 192mm, in-plane GRAPPA (R = 2), echo spacing = 0.68ms. The protocol is optimized to view mPFC in high-resolution and therefore the partial volume slab cuts off portions of the motor cortex and parietal lobe. EPI-based fieldmaps of positive and negative polarity were also collected before each run with similar parameters as the sequence.

#### Standard resolution data

A wholebrain multiband echo-planar imaging (EPI) protocol was collected in eleven participants with a 2.5 mm isotropic voxel size and the following parameters: multiband acceleration = 4, 72 slices, TR = 1120ms, TE = 30ms, A-P phase encoding, −30 degrees slice orientation from AC-PC line, flip angle = 54°, FOV= 192mm x 192mm. EPI-based fieldmaps of positive and negative polarity were also collected before each run with similar parameters as the sequence.

### fMRI Preprocessing

Data was preprocessed using a standard pipeline for preprocessing of multiband data. Using FSL (Smith et al., 2004), images were brain extracted, realigned, high-pass filtered (100s threshold), and unwarped. Images were denoised by ICA component removal. Components were extracted using FSL’s Melodic, classified into signal or noise with a classifier trained on separate datasets or manually classified for the high-resolution dataset. T2 images were aligned to T1 images with FSL FLIRT, and then both were normalized to standard space using ANTs (using CIT168 high resolution T1 and T2 templates (Avants et al., 2009; Tyszka and Pauli, 2016)). The functional data was first co-registered to anatomical images using FSL’s FLIRT, then registered to the normalized T2 using ANTs. For univariate analyses, data was spatially smoothed in FSL with a 5-mm FWHM Gaussian kernel. For multivariate analyses, data was spatially smoothed with a 2-mm FWHM Gaussian kernel.

### Predictions of Normalization

Let the vector 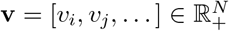 denote the values of individual items 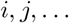 that make up a bundle. We derive formal results about the normalization function 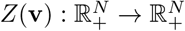

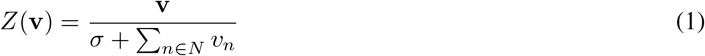

with contrast parameter *σ >* 0, which implements a linear constraint on neural activity (Webb et al., 2020). For simplicity we will derive these results for two-item bundles, but the argument extends easily to *N* -item bundles.

#### Concavity of Normalized Valuations

Let *v*_*i,j*_ denote the value of the bundle of two items *i* and *j*. Theories of linear value integration would imply that *v*_*i,j*_ = *v*_*i*_ + *v*_*j*_. However, if value is constructed via a normalization function (1), and the valuation of a bundle is the sum of the normalized valuations, 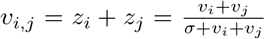, this relationship is concave in the sum of the item values when *σ >* 0 (see Figure 5).

**Figure 5:**
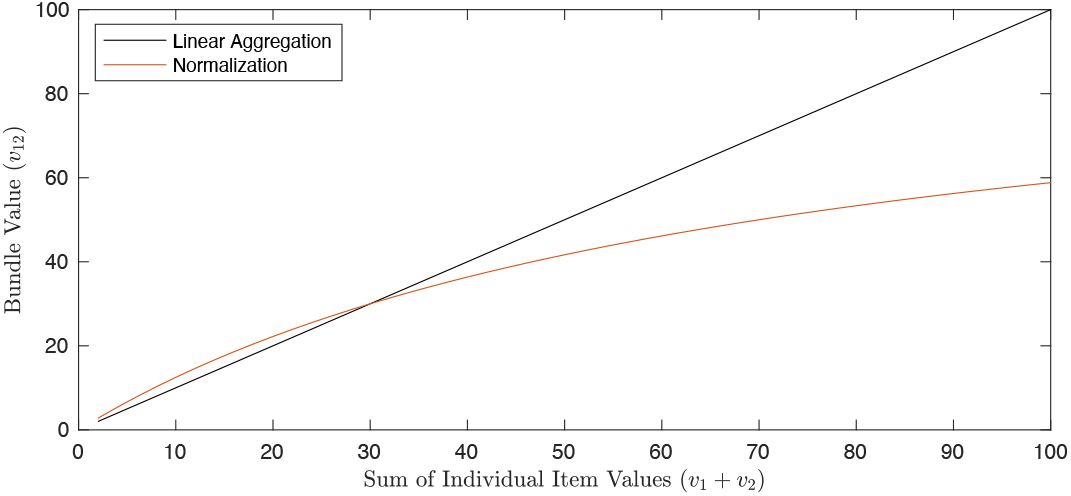
A normalization model predicts that the bundle value should be a concave function of item values if the reported bid is the sum of the normalized values.

**Figure 6:**
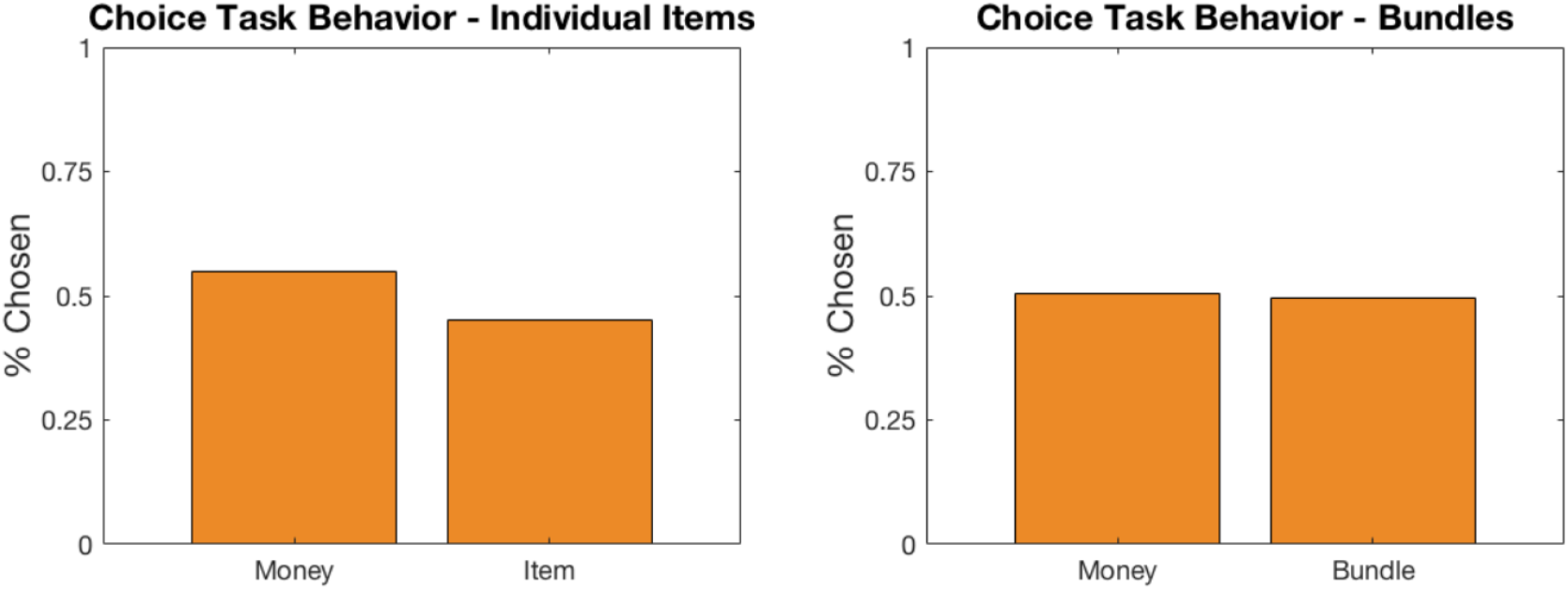
Supplementary Figure 2. Behavior on the choice task. Percentage of trials in which the item or bundle was chosen vs. the reference monetary amount.

**Figure 7:**
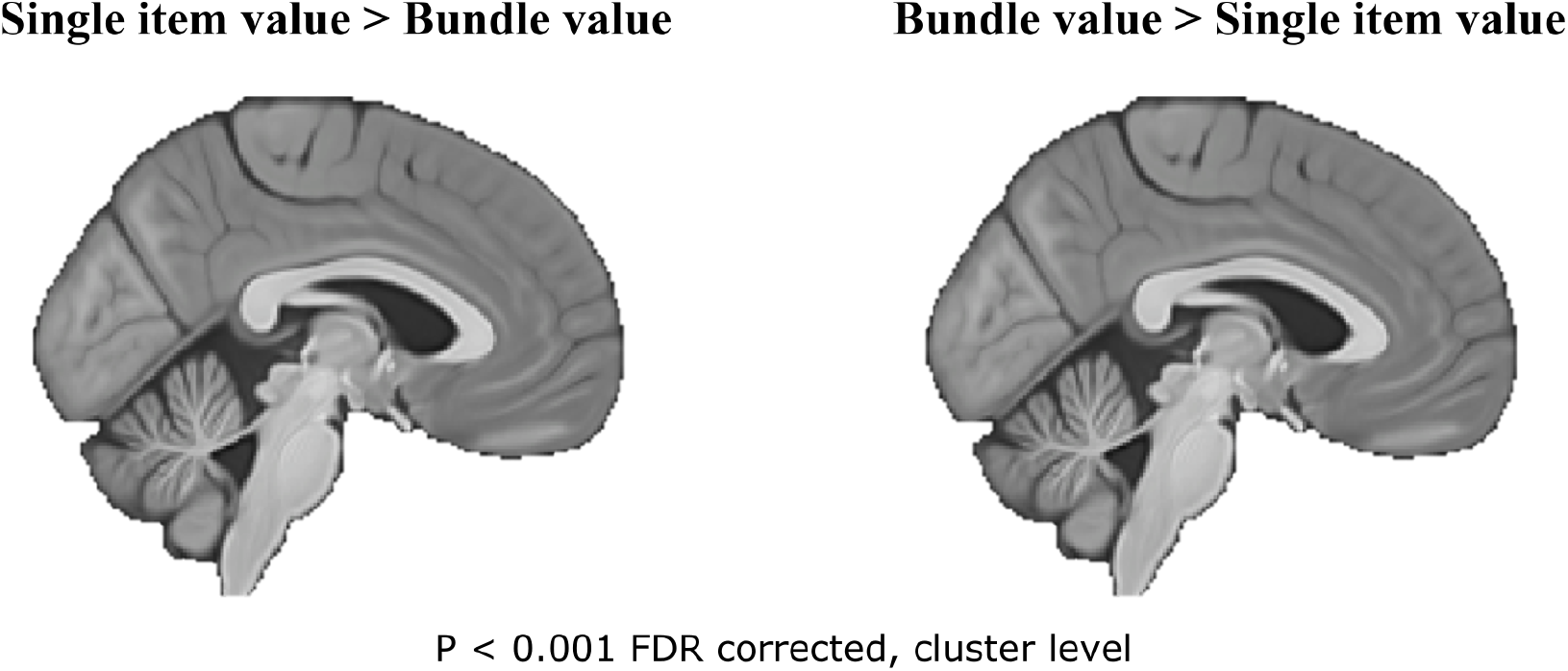
Supplementary Figure 3. Bundle Value vs Single Item Value. Univariate contrasts testing the interaction of value and trial type. No clusters survived in either comparison after multiple comparisons correction.

**Figure 8:**
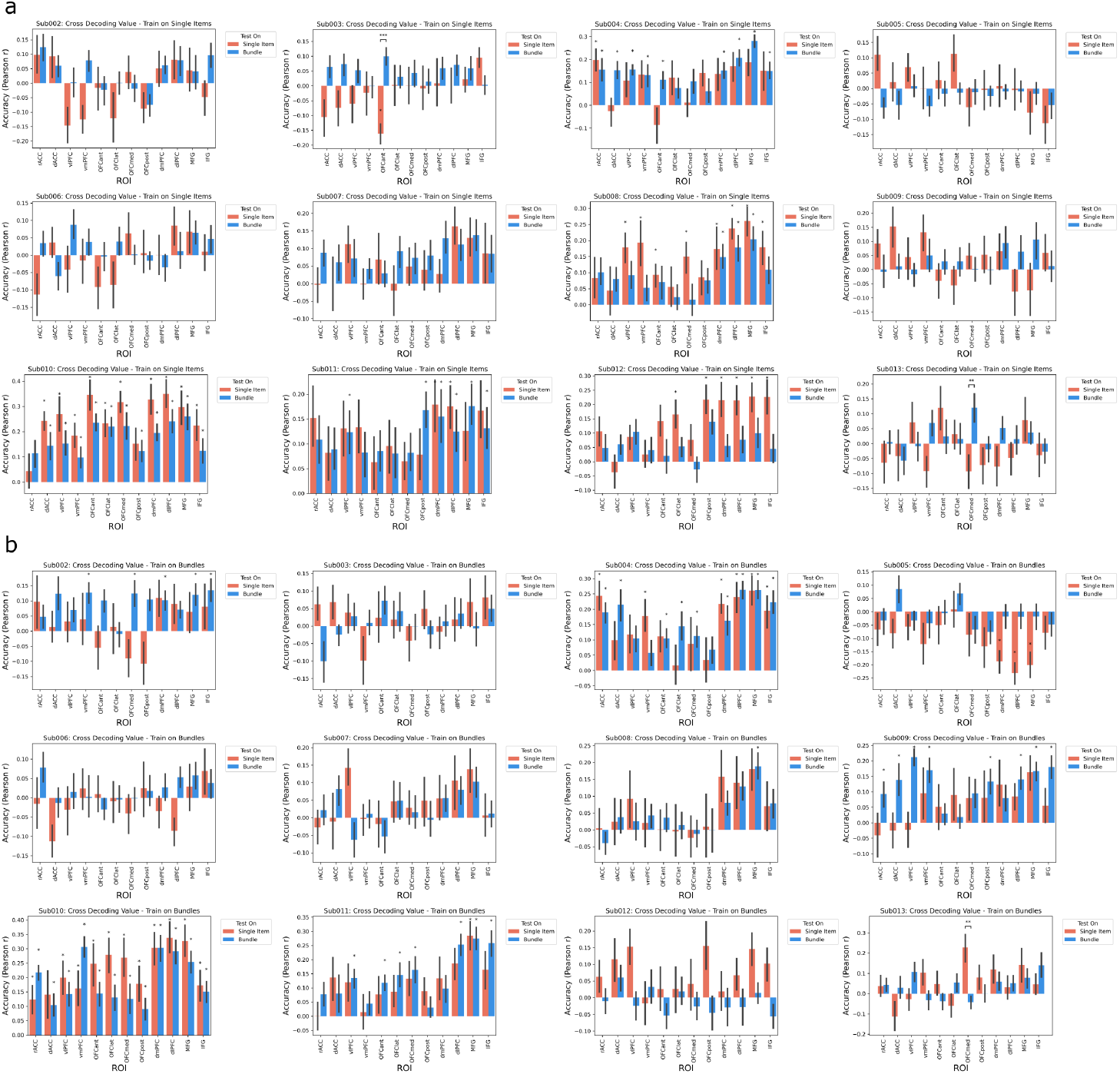
Supplementary Figure 4. Bundle Value vs Single Item Value. MVPA cross decoding analysis individual subject results. Left: decoders trained on trials of single items. Right: decoders trained on bundle trials. Asterisks * represent significant prediction accuracies on a test partition (two-sided one-sample Wilcoxon signed rank test P *<* 0.05 and FDR-corrected for multiple comparisons q = 0.05). At the group level, there were no significant paired differences in prediction accuracies between test conditions for any ROI (two-sided two-sample Wilcoxon signed rank test P *<* 0.05 and FDR-corrected for multiple comparisons q = 0.05). Error bars reflect SE across participants.

*Proof*. Let 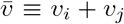. Therefore the bundle valuations can be written as a function of this sum, 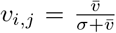. The second derivative of this function with respect to that sum, is 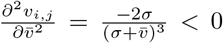, since *σ >* 0. Therefore it is concave.

#### Attenuation of Normalized Value for Bundles

The responsiveness of the value normalization function, with respect to one or all of its arguments, will be reflected in its first derivative. If the measurement of neural activity is linear in value, then this derivative will determine how neural activity responds to a change in value. We show here that for a normalization given by equation (1), the neural measure will attenuate for bundles. For simplicity, we consider the derivative of (1) with respect to an arbitrary argument *v*_2_,

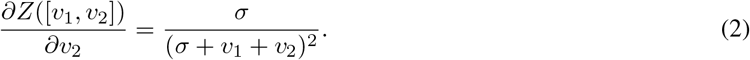

This derivative is positive and strictly decreasing in *v*_2_. In particular, when *v*_2_ = 0, and the normalization function is applied to only a single item, the first derivative of the function (i.e. its responsiveness) will be largest. It will therefore be attenuated for bundles (*v*_2_ *>* 0). If the measurement of neural activity of a bundle is linear in these normalized values, 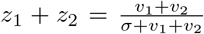, then its derivative is the sum of (2) for both items and the responsiveness of this measurement to value would be similarly attenuated on bundle trials. Intuitively, the responsiveness of the normalization function attenuates because there is an additional item that must be normalized against (both the numerator and the denominator increase by the same amount).

### Behavioral Analyses

Linear and nonlinear regression analyses were performed to model how bundle value is computed as a function of the values of the constituent items in the bundle, *v*_*ij*_ = *f* (*v*_*i*_, *v*_*j*_). The linear model represents bundle value as a linear combination of the individual item values:

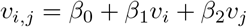

Item *i* and item *j* simply correspond to the item shown on the left and right, respectively, during stimulus presentation. A mixed-effects model was estimated across all subjects, with subject-specific random effects terms for intercept and slope. Additional nonlinear mixed effects models were also constructed.

Power:

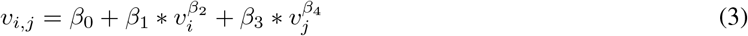

Logarithmic:

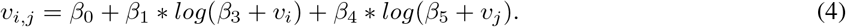

Normalization:

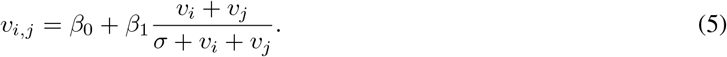

All models and model statistics were estimated with Matlab, with the *fitlme* and *nlmefit* functions, and model fits were evaluated with BIC and R^2^. Reported BIC scores reflect the sum of BIC across participants. To plot the fitted line/curve for each model in Figure 2.b, bundle value was computed from the fitted function parameters with *v*_*i*_ set equal to *v*_*j*_ for all values from 0-10 (with a 0.01 step size). The sum of these values (double of the value set both for *v*_1_ and *v*_2_) is represented by the x-axis.

The linear models were also separately fit on data from each type of bundle, food, trinket, mixed, and duplicates of the same item. Due to a small amount of data per subject, random effects terms could not be properly estimated for bundles of the same item, and therefore only a fixed effects model was estimated.

### Regions of interest

Regions of interest (ROIs) were defined using the AAL database (Tzourio-Mazoyer et al., 2002). The labels used in the paper are mapped to the original ROI names as follows: ACC pre: rACC, ACC sup: dACC, Frontal Inf Orb 2: vlPFC, Frontal Med Orb: vmPFC,OFCant: OFCant, OFClat: OFClat, OFCmed: OFCmed, OFCpost: OFCpost, Frontal Sup Medial: dmPFC, Frontal Sup 2: dlPFC, Frontal Mid 2: MFG, Frontal Inf Tri: IFG.

### Univariate Analyses

Univariate analyses were conducted in SPM12. General linear models (GLMs) were constructed to examine how subjective value and choice are encoded in the brain during the choice task. This GLM included a regressor for value, time locked to the onset of the trial as a parametric modulator, which was modified in separate GLMs to test hypotheses about value representation. The stimulus onset regressor additionally had another parametric modulator for trial type (−1 for single item trials and 1 for bundle trials). Thus, a contrast corresponding to the representation of bundle value or single item value could be computed as the interaction between value and trial type. Reaction times were additionally included as a third parametric modulator. Regressor for the choice made on a trial was time locked to the time of choice, with separate regressors for choosing an item/bundle and choosing the reference monetary amount. Regressors of no interest were included: left and right button presses (duration=0), motion regressors, and run. Missed trials were not modeled. Data from all three experimental days for each subject were consolidated into a single model, with separate sessions and distinct nuisance regressors for each day. The three participants that underwent the high-resolution scanning protocol are absent from the univariate analysis because that protocol was focused on ventral and frontal areas (with a limited field of view) with no coverage in dorsal frontal areas.

Different representations of value were included in the value regressor. The value contrast in Figure 4b,c used a normalized version of value, where value was z-scored by trial-type (item or bundle). The absolute value model uses the raw WTP $ amount that the subject rated an item/bundle outside the scanner. WTP minus reference (value difference) used WTP minus the reference monetary amount on that trial.

### Multivariate Analyses (MVPA)

To examine the nature of the bundle value code as the fine-grained distributed level, we implemented MVPA and RSA analyses. All analyses used the PyMVPA toolbox (Hanke et al., 2009), Scikit-learn functions, and custom Python code.

#### MVPA samples

To prepare for MVPA and RSA, we extracted trial-by-trial voxel-wise fMRI responses. A GLM was designed for each participant that modeled the onset and duration of each trial separately to extract the voxel responses that were unique to each trial. Other regressors of no interest modeled the other events and were not separated by trial (one regressor across an entire day): outcome phase (onset and duration), left and right button presses (duration=0). As in the univariate analyses, data across all three days in a subject were included in the same model, with different days and separate regressors per day entered as different sessions. After estimating the models, the parameter estimate maps (beta maps) for each trial were concatenated into a 4D dataset, with length equal to the number of trials a subject completed across the experiment.

#### Cross-Decoding Analysis

To test for distinct bundle value codes, single item value codes, and general value regions, we implemented a cross-decoding analysis similar to previously used methods (McNamee et al., 2013) at the ROI level. The 4D dataset of beta maps for every trial were loaded with PyMVPA functions. Value (the WTP bid) was z-scored by trial type (item or bundle) and included as the targets to predict in the PyMVPA dataset. Then the voxels for each ROI were entered as features for the cross decoding procedure. Ridge regression was used as the decoder in all analyses (Scikit-learn’s, linear model.Ridge), with alpha = 10^3^ (this parameter was optimized with sweeping). Each run was used as a separate cross-validation fold. Decoders were trained on the training samples from one trial type and tested on samples from the held out run (leave-one-run-out cross-validation). Decoders were tested on the samples from both trial types in the test set, even though they were trained on one type. However, decoder predictions were quantified separately for individual item samples in the test set and bundle samples in the test set. This ensured that performance could be compared across trial type separately from within trial type. Prediction accuracy was quantified by the Pearson correlation between the predictions and true value labels and averaged across cross-validation folds. This resulted in four decoder prediction accuracy metrics for each ROI and subject according to the four train/test splits: ‘train and test on single items,’ ‘train and test on bundles,’ ‘train on single items/test on bundles,’ ‘train on bundles/test on single items’. The average prediction accuracy across participants for each of these train/test splits is plotted in Figure 3c. Significance was assessed for each split vs. chance level (r=0) with two-sided one-sample Wilcoxon signed rank tests at *p <* 0.05 and FDR-corrected for multiple comparisons (ROIs) at *q* = 0.05. To assess if a decoder’s prediction accuracy was significantly different between conditions in the test set, we used two-sided two-sample Wilcoxon signed rank tests at *p <* 0.05 and FDR-corrected for multiple comparisons (ROIs) at *q* = 0.05.

#### Representational Similarity Analysis (RSA)

To examine whether the representational geometry of the regions of interest correlated to an absolute value or relative value code, we conducted representational similarity analysis (RSA). The 4D dataset of beta maps for every trial were loaded with PyMVPA functions as in the cross decoding analysis. Trial by trial neural dissimilarity matrices (DSMs) were constructed for every ROI by computing pairwise comparisons of the beta map across trials with PyMVPA’s PDist function. Euclidean distance was used as the distance metric. Two model DSMs were constructed: 1. trial by trial pairwise distances according to the difference in absolute $ value of the stimuli between trials 2. trial by trial pairwise distances according to the difference in normalized value of the stimuli between trials (where value was z-scored by trial type (item or bundle)). For all DSMs, within day comparisons were removed to avoid potential confounds due to similarity being driven by patterns being in the same run or day. Neural DSMs and model DSMs were then compared with Pearson correlations.

These neural DSMs were then compared against several theoretical model DSMs representing different hypotheses about value encoding:

##### 1. Absolute Value Model

Value is represented according to raw WTP bids, regardless of trial type (individual item or bundle).

##### 2. Subtractive Normalization Model

Value representation is normalized by subtracting the mean value within each stimulus category, adjusting for category-specific baselines.

##### 3. Z-score Normalization Model

Value is normalized within each trial type using z-scoring (subtracting the mean and dividing by the standard deviation), allowing for direct comparison between individual items and bundles despite their different value distributions.

##### 4. Full Normalization Model

Value is scaled using a biologically-plausible divisive normalization process, where the value coefficient can vary between trial types and includes a normalization factor that adapts to the local context.

##### 5. Null Model

Contains only nuisance regressors that account for potential confounds, including left vs. right button presses and item or bundle trial to account for the large visual differences between the trial types.

#### Full Normalization Implementation

The full normalization model implemented a context-dependent value scaling mechanism that accounts for trial-type specific adaptations in the neural value code. For each trial type (individual items and bundles), the model computed normalized values as:

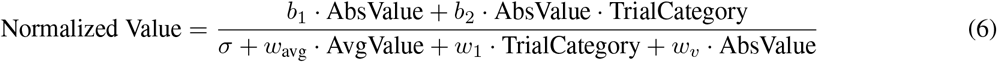

Where:

- AbsValue is the raw WTP bid amount
- TrialCategory is a binary indicator of trial type (individual item vs. bundle)
- AvgValue is the mean value within each trial type
- *b*_1_, *b*_2_, *σ, w*_avg_, *w*_1_, and *w*_*v*_ are fitted parameters

This formulation allows the model to:

1. Scale values differently for individual items versus bundles
2. Implement a non-linear compression of the value representation
3. Adjust the normalization factor based on the overall distribution of values in each context

The model implementation computed pairwise distances between these normalized values to create a model DSM. This was then combined with nuisance regressors through the formula:

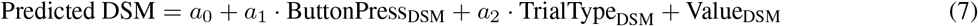

Where *a*_0_ through *a*_2_ are fitted parameters that control the contribution of each regressor to the predicted neural representation.

All models were fit to the neural DSMs using non-linear least squares optimization implemented in the LMfit Python package. To avoid potential confounds from within-day temporal autocorrelations, comparisons between trials from different scanning days were prioritized in the analysis. The goodness-of-fit for each model was assessed using Bayesian Information Criterion to assess the accuracy between the model predictions and the empirical neural DSMs while correcting for the number of parameters in the model.

**Table 1:**
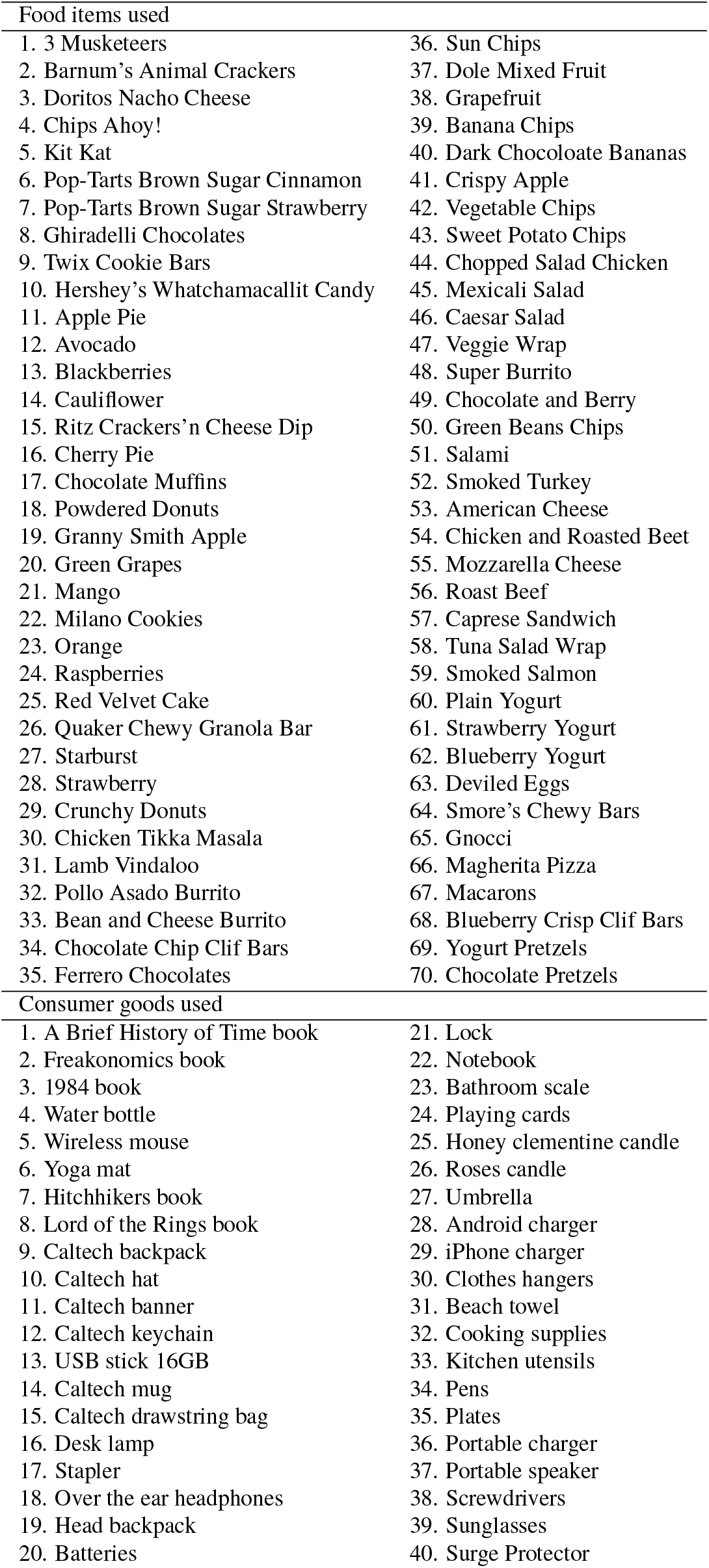
Items used in experiment.

